# *Sapodilla* rTLP exists as a monomer, dimer with β-1, 3-glucanase and antifungal activity

**DOI:** 10.1101/2023.03.22.533889

**Authors:** Chandana Thimme Gowda, Kammara Rajagopal

## Abstract

The present study deals in understanding the structure-function relationship of *Sapodilla* thaumatin-like protein (TLP). Most of the TLPs known to be stimulated in response to biotic, and abiotic stress. Few TLPs possess both antifungal and enzymatic properties, only very few TLPs possess either of the activity or none of the attributes. This characteristic of TLPs offer great challenges to examine its functional differences among its members, though they are structurally homologous. Therefore, we were concerned to see the functionality of *Sapodilla* TLP, by cloning in *E*. *coli*, expression, purification, and characterization. Being a plant derived protein, it possesses post-translational modifications such as the presence of disulfide bonds. Hence, we proposed to adapt various protein purification tools to purify and to obtain biologically active protein. The refolded and purified rTLP (recombinant TLP) exists as a monomer and dimer with β-1, 3-glucanase, and antifungal activity. The structure, function, relationship studies of rTLP (through deletion and site directed mutagenesis), observed to knock out the dimeric nature. Lastly, structural bioinformatics of rTLP reveal that their primary structural types are - β- and non-helical structures.

## Introduction

Plants have developed their sophisticated innate immunity to neutralize the effector molecules of pathogens. Concurrently, various pathogens, have been building up the immunity to counter attacking effector molecules and preserve their pathogenicity resulting in the pathological condition. Many plant pathogens have reported to impair plant development and growth [1]. Plants recognize the pathogenic elicitors that trigger the immune system through two modes beginning with the pattern-triggered immunity and the other effector-triggered immunity. This indicates the engagement of the signalling machinery to evade the pathogen infections [2]. Plants have evolved many strategies and the signaling pathways based on the inputs by pathogens at different checkpoints in the circuit of plant resistance. The elucidation of mechanisms beneath the signalling network has inspired researchers to realize more about the operation of plant immunity in response to specific pathogenic attacks.

### Pathogenesis-related proteins

PR proteins reported to accumulate in tobacco leaves as a hypersensitive response to the tobacco-mosaic virus infection [3]. PR proteins accumulated not just at the site of infection, but also systemically induced in plants. This, in turn reflects on the correlation of hypersensitivity response to systemic acquired resistance which sets up the host to regress further invasion by pathogenic microbes [4].

### Pathogenesis-related proteins in fungal pathogenesis

Pathogenesis-related proteins are a novel class of proteins possessing different enzymatic activities induced as a vital part of incompatible interactions in the plant pathogens and thereby resulting in the impediment of further pathogen invasion [5]. The process of inducible defence strategy begins when plants identify the microbes-secreted effector molecules and the wares of the R genes bind to these effectors [6], or the microbe, or pathogen-associated molecular patterns are picked out by plant pattern recognition-receptors [7]. Apart from transcription factors, ER-resident proteins, DNA repairing-proteins, these inducible PR proteins play a vital function in systemic acquired resistance [8] in combating fungi. They show strong induction in response to the biotic stress factors, thus they are recognized as ‘PR proteins’.

### Thaumatin

Thaumatins are intensely sweet-tasting proteins encoded by the palatability genes. They have similar sensory properties to those of three other types of sweet proteins: monellin, curculin, and miraculin [9]. The minimum concentration required to register the sweet taste is about 10^-8^ M, similar to hormone receptor interactions [10]. The retention of their stability at extreme temperatures and pH, and also its overall functional attributes suggest that, they are used to create different foods and beverages [11]. TLP’s (Thaumatin-like proteins) are monomeric proteins having a single polypeptide chain of approximately 200-207 amino acid residues; the most abundant amino acids are glycine, proline, cysteine and threonine. It is evident that TLPs possess a thaumatin signature sequence represented by -G-X-[G/F] -X-C-X-T-[G/A] -D-C-X (1,2) -[G/Q] -X (2,3) -C- the stretch of sequence [12] which appears in the mature polypeptide of the protein. The N-terminus amino acid sequence and the C-terminal extensions are highly conserved among the members of this family [13].

TLPs do possess β-glucan binding property, β-1, 3-glucanases are widely disseminated and are assorted into a different class of pathogenesis-related proteins (PR-2 family). Yet, unravelling a new facet of thaumatin-like proteins as glucan hydrolyzing enzymes. The enzymatic activity of TLPs has now been nominated to be one of the means of preventing pathogenic propagation into the plant tissues [14]. The presence of electronegative cleft of the protein has been suggested to be the key standard for the pronounced enzymatic activity of TLPs.

## Materials and Methods

Common Bio-chemicals, molecular biology reagents/salts, antibiotics, substrates were sourced from Hi-Media (Mumbai, India), Sigma-Aldrich (St. Louis, MO, USA), Sisco Research Laboratory (SRL, Mumbai, India) Invitrogen and MBI-Fermentas. The microbiological media like Nutrient agar, Luria-Bertani (LB) broth/media, LB Agar, Antifungal assay media (Agar), Potato Dextrose Agar were procured from Hi-Media (Mumbai, India) / Diffco were used for growth and maintenance of *E. coli* and fungal strains, respectively. *E. coli* cloning host XL1BL (Invitrogen, CA, USA), expression host BL21 (DE3) and the pET system were obtained from Novagen, Madison, USA.

### Gene synthesis and Cloning

The gene sequence of the *Sapodilla* acidic TLP encoding TLP protein was optimized for *E. coli* expression using M-fold analysis [15] and *E. coli* codon bias analyser online tools. The partial sequence of the TLP gene deposited in the GenBank sequence repository (Accession no. JN624813.1) was modified to optimize it for efficient expression in the heterologous system. The obtained composite gene was sub-cloned into the pUC57 a cloning vector. The resulting pUC57-TLP gene construct was transformed into *E. coli* XL1BL strain.

### Amplification of TLP and variants

The primers were designed specifically for the N- and C-terminal regions of the *Sapodilla* acidic TLP gene. The chemically synthesized TLP gene cloned into the pUC57 vector was used as a template. The gene was amplified using specific primers (mentioned in Table 1 using) both forward and reverse primers containing restriction sites for Nco I and Xho I. The PCR cycling parameters were programmed as follows; 3 minutes of initial denaturation followed by 35 cycles of 94 °C for 30 seconds, 52 °C for 45 seconds and 72 °C for 45 second amplification and a final extension of 5 minutes at 72 °C. The resulting product was analysed through 1.0 % agarose gel electrophoresis and further purified by using the GeneJet gel extraction kit.

**Table 1:**
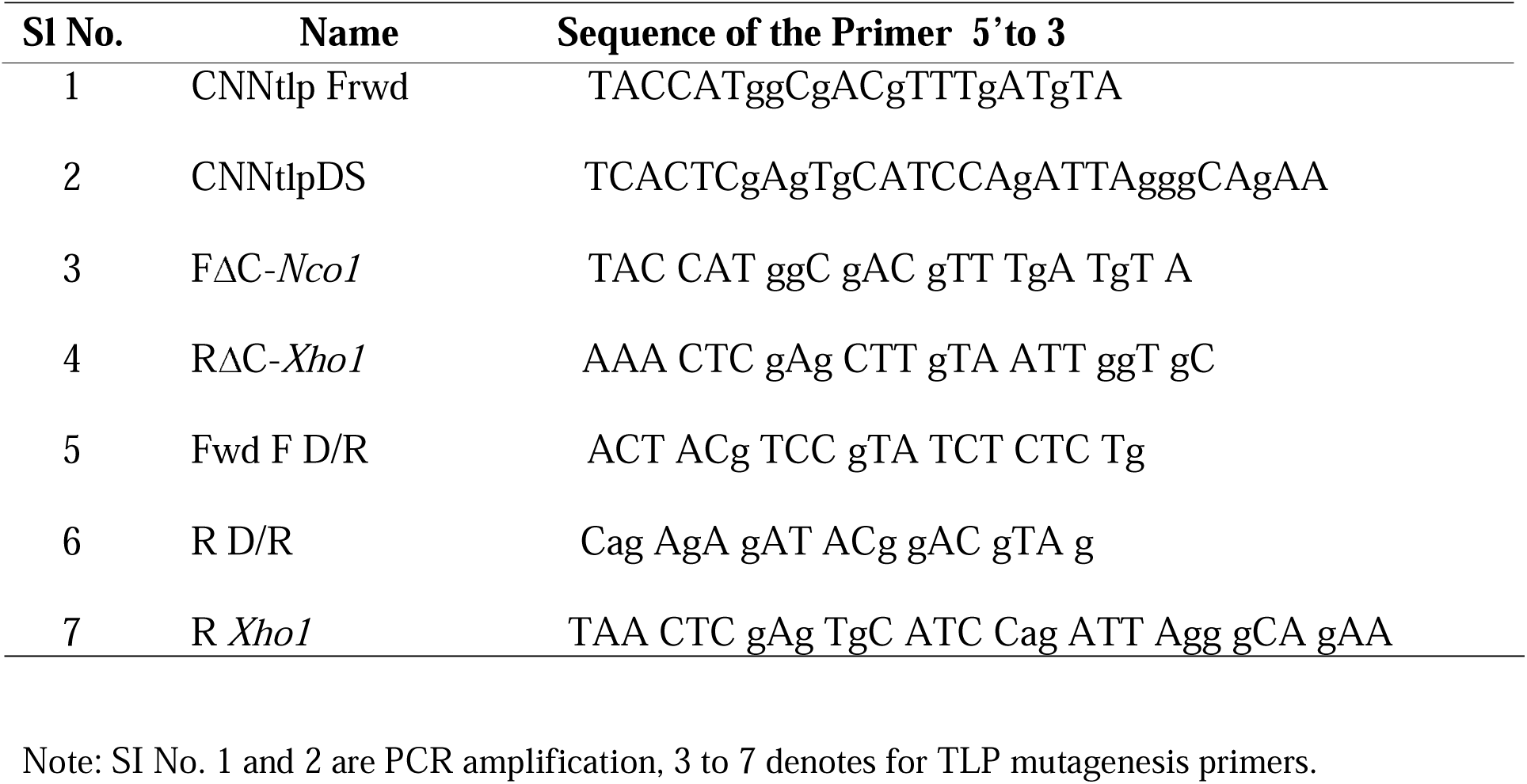
Primers used in this study

**Table 2:**
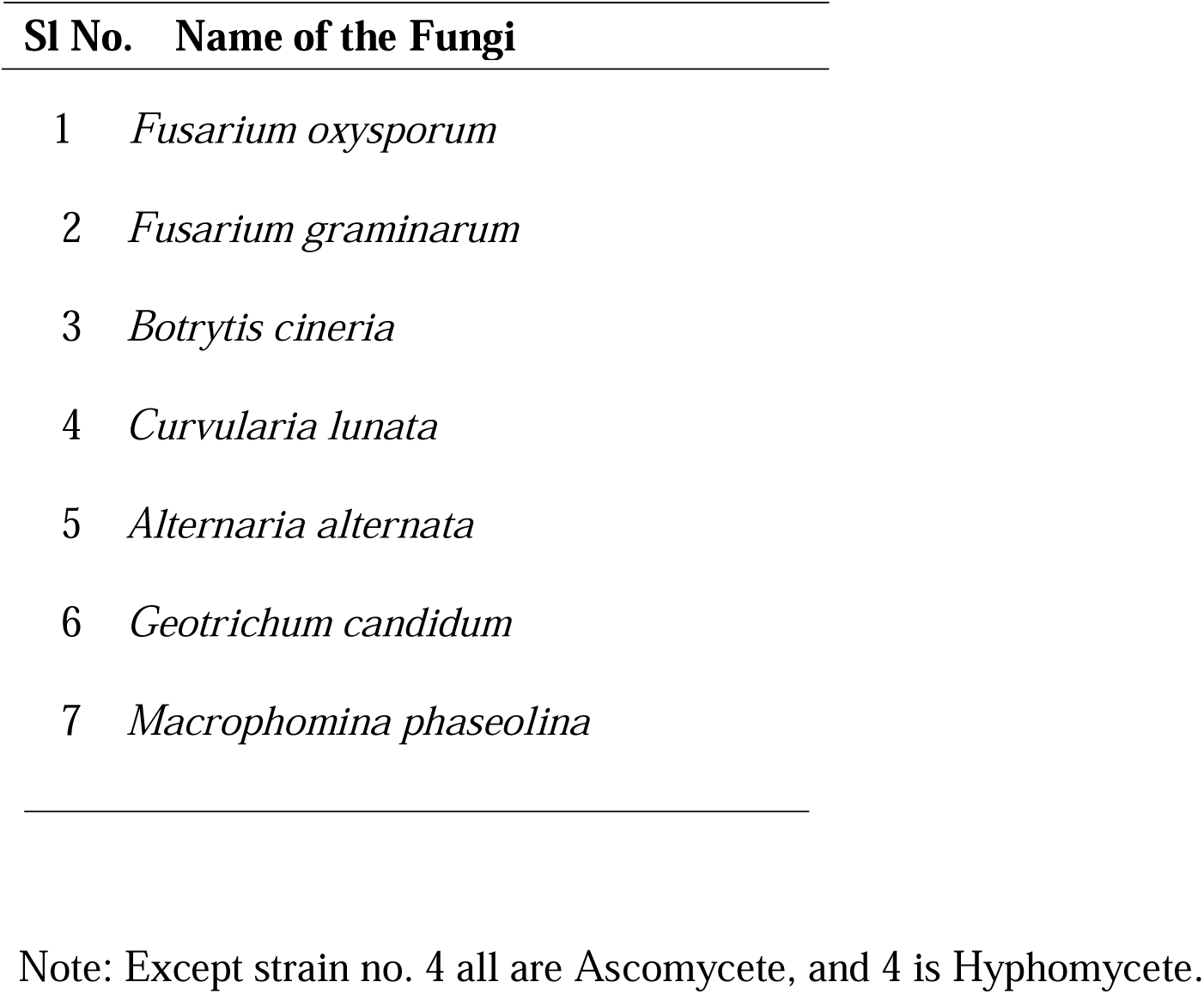
Fungal strains used in the study

| Sl No. | Name of the Fungi |
| --- | --- |
| 1 | *Fusarium oxysporum* |
| 2 | *Fusarium graminearum* |
| 3 | *Botrytis cineria* |
| 4 | *Curvularia lunata* |
| 5 | *Alternaria alternata* |
| 6 | *Geotrichum candidum* |
| 7 | *Macrophomina phaseolina* |

Note: Except strain no. 4 all are Ascomycete, and 4 is Hyphomycete.

### Construction of rTLP and various mutants in pET23 (d) plasmid

The purified TLP gene flanked by Nco I and Xho I (Fermentas Inc.) restriction sites was sequentially digested by FastDigest Nco I and Xho I restriction enzymes. The vector pET- 23d ^(+)^ was also sequentially digested with FastDigest Nco 1 and Xho 1 restriction enzymes. The double-digested linear vector was excised and purified. The digested, purified TLP gene (insert) and pET-23d ^(+)^ (vector) were quantified by NanoDrop ND-1000 spectrophotometer. Later, the purified vector was dephosphorylated by CIAP (Calf intestinal alkaline phosphatase, NEB). The insert and the vector were ligated at a 5:1 molar ratio. The ligated products were purified and transformed into competent *E. coli* XL1BL.

### Construction of rTLP mutants

Two variants of rTLP were constructed (i) a truncated protein by deleting C-terminal nonapeptide (-VVFCPNLDA) (ii) Mutant obtained by replacing aspartic acid at 96^th^ position by arginine residue. The C-terminal deleted truncated protein was obtained by the conventional PCR method using the primers listed in Table 1. The site-directed mutagenesis was carried out by Splice Overlap Extension PCR (SOE-PCR) [20] method.

### Induction, Expression of rTLP and its variants

The selected recombinant constructs harbouring the desired gene were transformed into *E. coli* BL21 (DE3) expression strain. A single isolated colony grown on LB agar substituted with 100 μg/ mL of ampicillin was selected and revived in the required amount of LB broth supplemented with ampicillin (100 μg/ mL) by growing overnight at 37 °C. Sub-culturing of 4.0 % of overnight grown inoculum was used to inoculate 1.0 litre of LB medium. The bacterial culture was grown till it reaches an OD_600_ of 0.60 by continuous shaking with 200 rpm at 37 °C. Subsequently, induced with 1.0 mM IPTG. The bacterial cultivation was carried on by continually shaking in an incubator shaker 200 rpm at 37 °C for 5 h. The cells were harvested by centrifugation at 5000 rpm for 15 min. They were re-suspended in 50 ml ST buffer and subjected to sonication for 15 min 30 secs off, 30 sec’s on the mode in a probe sonicator. The cell lysate obtained was analysed for recombinant protein by SDS-PAGE electrophoresis and subsequently, subjected to various methods of purification. The identical induction, expression, and purification procedures were followed for various rTLP variants.

### Preparation of Inclusion Bodies (IBs)

After sonication, the cells were centrifuged the supernatant and the pellet were separated. The cell pellet containing the IBs was washed thrice with ST buffer (50 mM Tris- Hcl containing 300 mM NaCl at pH 8.0 substituted with 0.1% of Triton X-100). The process rendered to obtain clarified IBs by greatly reducing other contaminating proteins.

### Solubilisation of Inclusion Bodies (IBs)

The IBs were solubilized using different concentrations of urea (in ST buffer) from the range of 2.0-8.0 M to determine the optimum urea concentration required for complete denaturation / solubilisation of IBs. The IBs were found to be completely solubilized by 8.0 M urea solution suspended in ST buffer. The mixture was continuously stirred for 3 h at room temperature to ensure complete denaturation and reduction of disulfide linkages in the proteins. The mixture was further clarified by centrifugation before loading onto the IMAC purification column.

### Refolding and purification of recombinant proteins using metal-ion chromatography

The solubilized protein sample was refolded and purified in three different methods to compare the efficacy for purification, refolding and obtaining better yields of the bioactive protein. The methods were mainly categorized as; refolding before affinity chromatography and On-column refolding during purification.

### Refolding before affinity chromatography

In this method, the refolding was performed with two different refolding buffers; (a) Dilution with ST buffer; (b) Dilution with ST buffer containing reduced and oxidized glutathione (GSH: GSSG).

#### (a) Refolding by air oxidation (by diluting with ST buffer)

The solubilized IBs with 8.0 M urea was centrifuged at 10000 rpm for ten minutes. The supernatant collected was flash diluted with 100 volumes of ST buffer by dropwise addition of 8.0 M urea suspension. The mixture was allowed for air oxidation in open beakers at 4.0 °C overnight. The refolded suspension was set aside to stand for 2 h at 4.0 °C to fully precipitate the unfolded protein intermediates. The refolded proteins were collected by centrifugation and the supernatant was loaded onto pre-equilibrated Ni-NTA agarose purification chromatography.

#### (b) Refolding by dilution with ST buffer (containing GSH: GSSG)

In this method, an aliquot of IBs solubilized in 8.0 M urea was subjected to air oxidation in the presence of redox pair. The supernatant obtained after solubilizing was flash diluted with ST buffer substituted with 4.0 mM reduced and 1.0 mM oxidized glutathione (GSH: GSSG at 10:1 ratio). The mixture was allowed to refold at 4.0 °C overnight and centrifuged. The supernatant obtained was further subjected to purification by affinity chromatography.

The purification of rTLP was achieved by immobilized metal-ion chromatography using Ni-NTA agarose beads (Invitrogen, Life Technologies, CA, USA). The refolded mixture was loaded directly onto Ni-NTA resin equilibrated with a pre-equilibrating buffer (0.05 mM Tris- Hcl, pH 8.0 containing 0.3 M NaCl) at a flow rate of 0.5 mL/min. The column was subsequently washed with a pre-equilibrating buffer containing 10 mM imidazole followed by 50 mM imidazole respectively. The rTLP was eluted with 300 mM imidazole dissolved in pre-equilibrating buffer and collected 1.0 mL fractions. The fractions obtained were analysed for purity and functional activity by SDS-PAGE electrophoresis and zymography. The fractions containing rTLP of desired purity were pooled and dialyzed against 20 mM Tris-Hcl buffer, pH 7.2 for desalting. The dialyzed sample was concentrated using the Eppendorf concentrators and stored at 4.0 °C until used.

### On-column refolding during affinity chromatography

The fusion protein obtained after clarification and denaturation of IBs was directly loaded onto the Ni-NTA agarose column equilibrated with a pre-equilibrating buffer containing 8.0 M urea. The sample was loaded with a flow rate of 0.5 mL/ min. The flow-through from the column was collected and reloaded onto the column to ensure the maximum binding of rTLP proteins. The column was washed thoroughly until the UV absorbance at 280 nm baseline was obtained. The gradual refolding of bound proteins was achieved by washing the resin with pre-equilibrating buffer containing decreasing linear gradient of urea from 8.0 – 0.0 M substituted with 10 mM reduced glutathione and 1.0 mM oxidized glutathione (10:1 ratio of glutathione redox pair). The column was again washed with equilibration buffer containing 10 mM and 50 mM imidazole sequentially. The refolded and bound proteins were eluted in the presence of equilibrating buffer containing 300 mM imidazole. The post purification procedures followed for rTLP were followed for variants also.

### In-gel assay for β-1, 3-glucanase activity of recombinant proteins

The rTLP was analysed for β-1, 3-glucanase activity by an in-gel assay using native/ non-denaturing SDS-PAGE gels [16]. The procedure in brief; Purified rTLP was subjected to SDS-PAGE. Subsequently, the gel was washed with water, treated with 0.05 M sodium acetate (pH 5.0) for 30 min, followed by incubation in the mixture of 0.05 M sodium acetate (pH 5.0) and 1.0 g Laminarin (Sigma-Aldrich) in a total volume of 75 mL at 40 °C for 2 h. Subsequently, the gel was soaked in a mixture of methanol: water: acetic acid (5:5:2 v/v) for 5 minutes and washed with water twice. Finally, to stain for β-1,3-glucanase, the gel was soaked in 300 mg of 2,3,5-triphenyl tetrazolium chloride (TTC) in 200 mL of 1.0 M NaOH in a boiling water bath for 10 min until red coloured bands appeared.

### Antifungal activity assay by disc diffusion

The refolded and purified recombinant rTLP was tested for its antifungal potential by the disc diffusion method as described [17]. A mycelial disk of 6.0 mm was excised from the revived and full-grown PDA plates and placed on fresh PDA plates, allowed to grow for three days at 27 °C, sterile conditions. The sterile disks of 3.0 mm were placed adjacent to the actively growing mycelia periphery. The rTLP at different concentrations (12.5 μg, 25 μg, 50 μg, and 100 μg) were appended to the discs and incubated at 27 °C for 48 h. Each experiment was performed in triplicates and repeated in three independent days to obtain the statistical significance. BSA (20 μg) and kanamycin (25 μg) were used as negative and positive controls.

### Immuno-detection of recombinant proteins

The Western blot was developed by Towbin et al., [18] is an analytical technique widely used to distinguish specific antigens in a tissue homogenate or cell lysate. The method involves an electrophoretic separation of proteins, transfer of the resolved proteins to a membrane and visualizing the specificity of proteins for the labelled antibodies of interest [19]. Here we followed the similar method by using rTLP antibodies on native PAGE.

### Sequence analysis and structural bioinformatics

#### Homology modelling of TLP

*In-silico* based structural and functional prediction provides greater insights into the molecular framework and mechanisms of proteins. A 3D-modelling of *Sapodilla* acidic TLP was performed by submitting the primary sequence to the I-TASSER server. I-TASSER (Iterative Threading ASSEmbly and Refinement) is an integrated platform that generates a three-dimensional model using the sequence-to-structure-to-function paradigm [21]. This tool selects structurally close templates from protein data bank by a fold-recognition technique called *threading*. The top hits of the structural fragments obtained were then reassembled using computational simulations. I-TASSER generates five template-based models. The best model is selected based on the percentage of coverage with the template, the C-score, and the TM-score of the generated model. The optimized molecular model was visualized using PyMol or UCSF Chimera online visualization programs.

#### Verification of 3D model

The substantiation of the 3D model is an essential step to analyse the reliability of the model for extending the results into further investigations. The model evaluation was performed by online programs like PROCHECK, Verify_3D, and ERRAT [22]. PROCHECK analyses the stereo-chemical quality of the model and Verify_3D determines the 3D structure of the model [23]. ERRAT differentiates correctly and incorrectly folded regions of protein structures based on characteristic atomic interactions [24]. The secondary structure, proportion of *Sapodilla* acidic TLP protein was predicted using PROFUNC package protein structure prediction server [25].

### Ramachandran plot for computer generated TLP

PROCHEK analysis was performed to derive Ramachandran plot. Ramachandran plot, was made to determine permitted torsional angles that can obtain insight into the structure of TLP peptides.

### PROMOTIF Scan

PROMOTIF Scan was followed to identify the presence of common structural motifs and also their positions in the predicted structure of TLP. That helps to unravel the secondary structure of a protein.

### Quality assessment of the predicted rTLP model and Verify 3D-plot

The plot is generated by comparing the 3D structure with the primary sequence of the protein. Subsequently, a profile is created based on the local environment of the residues.

## Results

### Design and optimization of synthetic *Sapodilla* acidic rTLP gene for hyper-expression

The N-terminal nucleotide sequence, including the Shine-Dalgarno sequence were subjected to the online codon optimization software. It was optimized for secondary structure predictions by M-fold analyser. The nucleotide base pairs were replaced to avoid highly stable intra-molecularly base-pairing structures. The final sequence arrived by following M-fold analysis is as follows: ***<u>AAG AAG GAG ATA TAC ATATG GCG ACG TTT GAT GTAGTA AAT CAA TGT ACT TTT.</u>*** The above sequence was assembled with the previously reported TLP gene fragment resulting in a complete gene sequence of 633 bp that encodes a mature acidic TLP.

It was noteworthy that 33 codons from the nucleotide sequence were distinctly different from the overall *E. coli* codon usage. These codons were replaced by the *E. coli* preferential synonymous codons that can maximize the likelihood of overexpression of proteins in *E. coli* host without altering the encoding protein residues. The above-optimized sequence was chemically synthesized with unique restriction sites of Nco I and Xho I flanking the N- and C-terminal ends. This synthetic gene was subsequently inserted into the pUC57 vector by sub-cloning. The recombinant plasmid DNA of the pUC57 containing the TLP gene was first transformed into *E. coli* XL1BL. The resulting rTLP was used as a template for the amplification of the TLP and its variants. Finally, the pET-23d ^(+)^-TLP and variants were transformed into *E. coli* BL21 (DE3) strain for expression studies.

### Induction and Expression of rTLP and variants

The cells containing pET-23d ^(+)^-TLP were grown at 37 °C in 10 mL LB broth containing 100 μg/mL ampicillin under vigorous agitation. The cells were induced with 1.0 mM IPTG at 0.4 OD, were harvested after 5 h and analysed for the presence of rTLP protein. The pelleted cells were re-suspended in 2xSDS-PAGE loading dye and separated by 12 % SDS-PAGE (Fig. 1A). The induced fraction was shown to contain hyper-expressed rTLP protein (Fig. 1A, Lane 2) whereas un-induced fraction also showed a faint band of the same molecular mass indicating the leaky expression owing to the use of BL21 (DE3) cells (Fig. 1A, Lane 3). The molecular mass of hyper-expressed protein was found to be ∼25 kDa which was consistent with the expected molecular mass of TLP protein under reducing condition.

**Figure 1A:**
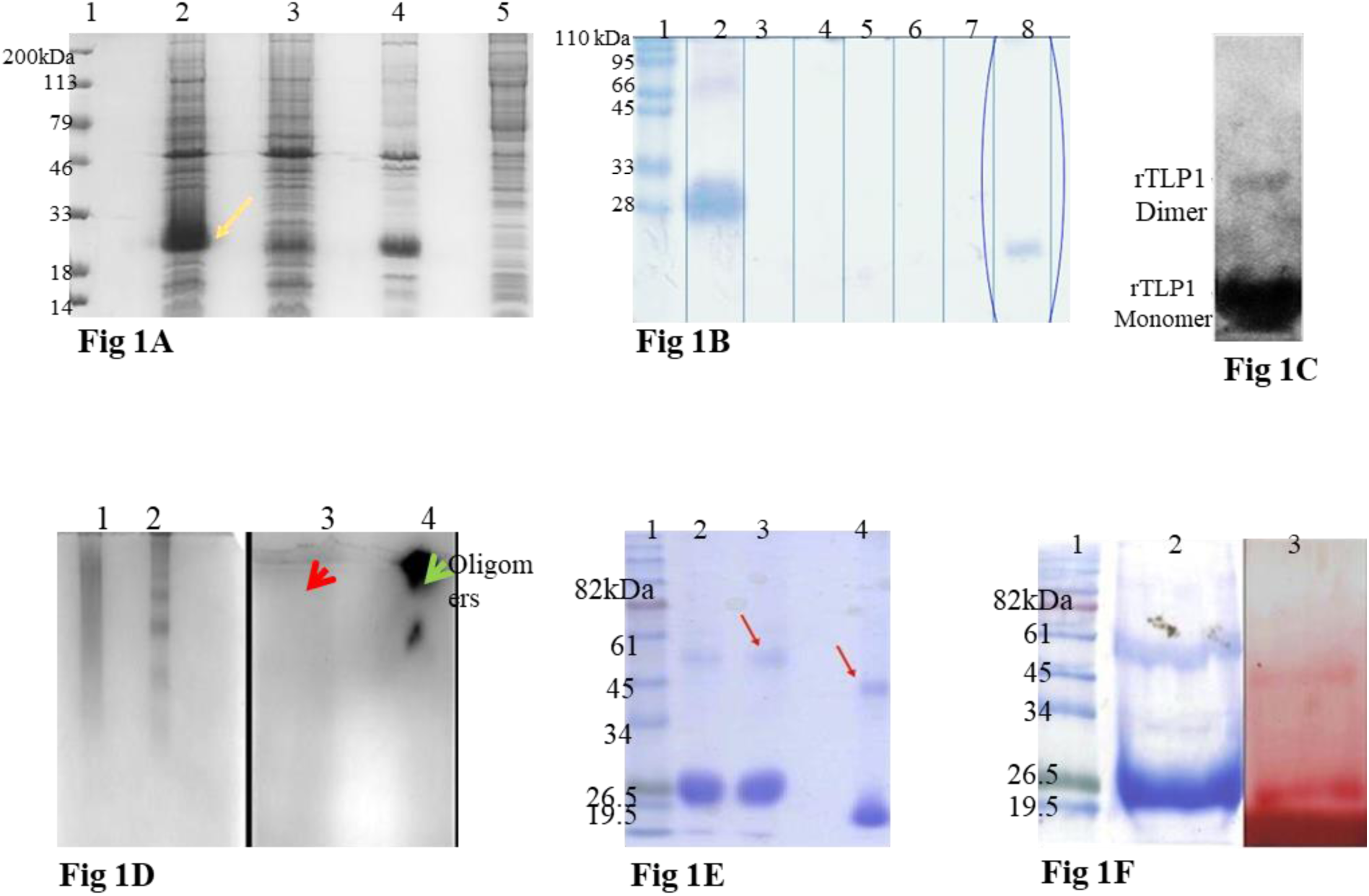
SDS-PAGE profile of overexpressed hexa-histidine tagged recombinant TLP protein. Lane 1: Protein molecular weight marker; Lane 2: pET-23d ^(+)^-TLP cell lysate induced; Lane 3: Cell lysate of un-induced, Lane 4: Sonicated pellet; Lane 5: Sonicated supernatant. Arrow indicates induced expressed protein TLP. **Figure. 1B:** SDS-PAGE electrophoresis profile of soluble TLP IBs. Lane 1: Protein marker; Lane 2: Insoluble inclusion bodies, Lane 3: Soluble fraction; Lane 4-7: Soluble fraction of IBs when washed with ST buffer containing 0 M, 2.0 M, 4.0 M and 6.0 M Urea respectively; Lane 8: Soluble fraction of 8.0 M Urea suspension. **Figure. 1C:** Western blot analysis of bacterially expressed rTLP using anti-TLP antibodies raised in rabbit. **Figure. 1D:** Native-PAGE electrophoresis profile of rTLP protein; Lane 1-2: rTLP, without β-ME; Lane 3-4: rTLP, with β-ME. Oligomers are shown in the gel. **Figure. 1E:** Determination of enzymatic activity of rTLP at native conditions. (A) Native-PAGE electrophoresis of rTLP with and without β-ME. (B) Zymography of rTLP with Laminarin as substrate. Red arrow indicates samples of β-ME treated, Green arrow indicates without. **Figure. 1F:** The enzymatic activity of rTLP as shown by zymography on non-reducing PAGE using laminarin as substrate. This figure indicates that the protein is active both in its dimeric and monomeric form.

### Solubility determination of rTLP

However, to determine the solubility of the expressed His_6_-tagged mature TLP protein, an aliquot of induced cell pellet was disrupted by sonication. The whole-cell lysate was centrifuged to separate the supernatant and pellet. The soluble fraction from the supernatant and the insoluble fraction from the cell pellet were separated on 12 % SDS-PAGE. It was observed that the sonicated pellet was shown to contain over-expressed protein, indicating that the rTLP protein is expressed as IBs) (Fig. 1A, Lane 4, and 5) and it is insoluble.

### Solubilization of rTLP Inclusion Bodies (IBs)

To determine the conditions for solubilizing the rTLP inclusion bodies, the IBs were serially diluted in ST buffer (50 mM Tris, 300 mM NaCl, pH 8.0), containing different urea concentration in the range of 2.0-8.0 M. The suspension was stirred at 4.0 °C for 15 min. and centrifuged, a portion of the supernatant was loaded onto 12 % SDS-PAGE. 8.0 M urea supernatant showed the presence of a band corresponding to the molecular weight of rTLP indicating that 8.0 M urea is the suitable concentration for solubilizing (Fig. 1B, Lane 8). No rTLP was observed in Lane 3 to 7, indicating these urea concentrations unable to dissolve IBs. The protein is discovered to be partially devoid of contaminants, which facilitated the preparation of the sample for easy purification.

### Immuno-detection of rTLP

Immunoblot experiment was performed to identify the presence of rTLP in the purified fractions. The specific polyclonal antibodies rose against native-sapodilla acidic TLP recognized the recombinant protein suggesting that the expressed protein product was TLP. A protein band corresponding to 45 kDa showed a positive reaction with the anti-TLP antibody revealing the presence of rTLP dimer (Fig. 1C). Two protein bands were observed that are placed one upon another indicating a monomer and dimer. The populations of monomer were considered to be much higher than the dimer population.

### *In-vitro* protein refolding and purification of rTLP

#### Dilution with ST (Sodium chloride and Tris.cl) / refolding buffer

In the first attempt, the rTLP concentration in 8.0 M urea suspension was adjusted to 5.0 mg/mL of solution. This suspension was flash diluted by adding into 10 volumes of ST buffer drop by drop, followed by air oxidation at 4.0 °C overnight. It was noted that there was a little precipitation of aggregated proteins, which was separated by low-speed centrifugation. The refolded protein was purified by metal-ion affinity chromatography, under the denaturing condition, the protein band corresponding to the molecular weight of 26 kDa was intense. There was another protein band corresponding to the molecular weight of 45 kDa, which was exactly double the molecular weight of rTLP (Fig. 1D). Figure. 1D Lane 1 shows a protein smear indicating intramolecular disulfide bonds. Therefore, to reduce them the protein was treated with β-mercapto ethanol (β-ME). After the treatment one can observe two major protein bands one at 26 kDa, and the other a dimer Lane 2. The rTLP treated with and without β-ME were subjected to western blotting results envisages oligomerization of rTLP (Fig. 1D, Lane 3, 4). A similar scenario of the presence of a dimer has been observed in the western-blot also (Fig. 1C). This protein band could likely be the dimer of the *Sapodilla* TLP protein. The protein yield obtained with this process before dialysis was found to be∼12 mg/L of bacterial culture; however, the yield of active protein was found to be much lower i.e., ∼3.0 mg/L of culture after dialysis. The extensive loss of protein by this method is attributed to the aggregation of proteins during refolding and dialysis. The cloudy precipitate in the suspension was observed after it was allowed to rest overnight at 4.0 °C.

#### Dilution with ST buffer containing GSH/GSSG redox pair

As thaumatin-like proteins are known to be disulfide-rich proteins, the formation of proper native disulfide bridges needs redox intermediates. It was shown in the previous experiment that mis-bridged disulfide species were responsible for forming oligomers with various degrees of oligomerization, and hence an alternative refolding buffer system was optimized. The denatured protein suspension was flash diluted into refolding buffer supplemented with reduced and oxidized glutathione at a ratio of 10:1, respectively. The protein suspension was left for refolding overnight upon continuous stirring at 4.0 °C.

### On-column refolding during affinity chromatography

Contrary to the above procedure, the protein refolding was performed during the column purification step. Since the protein contains poly-histidine tag, it is compliant to refolding on-column and offers a single capture step on immobilized metal ion chromatography. The on-column refolding of 8.0 M urea-denatured protein suspension was performed with linear gradient wash from 8.0 - 0 M urea containing 10:1 ratio of reduced and oxidized glutathione. The soluble protein was eluted with 0.30 M imidazole.

### Validation and analysis of refolded protein

Native-PAGE was performed both in the presence and absence of β-ME (Fig. 1E, Lane 1, 2). The recombinant protein showed a weak β-1, 3-glucanase activity which could be solely detected by zymography (Fig. 1E, Lane 3, 4). The protein was examined for its activity both in the presence and absence of β-ME. The untreated protein sample displayed endoglucan hydrolyzing activity by the release of oligosaccharides. This indicates that the protein is active both in its monomeric and dimeric forms. This also emphasizes on the essentiality of the proper folding and formation of disulfide linkages for the molecule to exhibit enzymatic activity.

It is demonstrated that the oligomers were broken down to monomeric molecules upon treatment with β-ME whereas multiple bands were seen in the untreated sample (Fig. 1E). Also, analysis by SDS-PAGE in denaturing condition showed the presence of dimeric protein (Fig. 1D). This attribute may be ascribed to the occurrence of intermolecular disulfide bonds due to increased protein concentration in the solution which may favour protein-protein interactions.

The native-PAGE analysis still showed the aggregated proteins stacked at the resolving-stacking gel interface. The protein fraction obtained after dialysis precipitated significant and do not preserve the functionality as demonstrated by zymography. Therefore, the protein yield after the purification process was found to be considerably poor.

The SDS-PAGE analysis of refolded and purified proteins under denaturing conditions shows a major band of 26 kDa indicating the presence of rTLP protein in monomeric form. In non-denaturing conditions (non-reducing, but denaturing), the change in electrophoretic mobility is observed wherein the band is shifted to 18 kDa molecular weight. As this protein has a high content of disulfide bridges that imparts its stability and compactness, this remarkable difference in electrophoretic mobility of rTLP suggests that the protein is compactly refolded (Fig. 1D). The appearance of a band corresponding to 45 kDa molecular weight (shown an arrow mark) suggests that dimer formation takes place both in denaturing and non-denaturing conditions (Fig. 1D, Lane 4). The dimer also exhibits a mobility shift which is a characteristic feature of rightly folded proteins with a high content of disulfide bonds like TLP’s. It is noteworthy that the aggregation of recombinant protein has been drastically reduced in this method as shown by a non-denaturing (non-reducing) protein profile. The yield of active and refolded protein was found to be ∼6.0 mg/L of bacterial culture.

The enzymatic activity of rTLP obtained by this method was elucidated by zymography on non-denaturing (non-reducing) polyacrylamide gel electrophoresis with laminarin as a substrate (Fig. 1F). Both monomer and dimer of the protein showed modest but detectable enzymatic activity on the substrate. The protein is observed to be active in its monomeric as well as dimeric forms.

### Evaluation of *in-vitro* antifungal activity by disc diffusion

Very common fungal pathogens have been chosen based on their pathological relevance to *Manilkara zapota*. Commonly, *Sapodilla* fruits are infected by the fungal pathogens during ripening and storage due to their soft texture and high sugar content [26]. They are known to be more susceptible to sour rot disease caused by *Geotrichum candidum* and black rot disease caused by *Aspergillus niger*. Likewise, soft rot disease caused by *Pestalotiopsis mangiferae* [27]. Similarly, *Rhizoctonia solani* and *Fusarium* spp are considered to be the most prevailing pathogens causing pinkish to red rots in the fruit. *Botrytis cinerea* is a common necrotrophic fungus causing grey rot and noble rot, especially in fruits and berries. Thus, the exploration of the antifungal property of rTLP could be of some relevance in plant disease management.

First of all, the antifungal activity of rTLP was found to be species-specific as demonstrated by disc diffusion assay. The sensitivity of rTLP was observed only for two pathogens, *Alternaria alternata* (Fig. 2A) and *Geotrichum candidum* (Fig. 2B). Even the highest concentration (100 μg) of the test protein did not affect the growth of other fungi. It is shown that at a concentration of 50 μg protein mycelia growth was only slightly inhibited (Fig. 2A, B arrows indicate the growth inhibition). Between the two species, the growth inhibition of *Alternaria* was relatively distinct from that of *Geotrichum candidum*. Also, *Geotrichum* spp. being a prolific pathogen causing serious disease conditions in *Sapodilla*, it could be one of the inducers of biotic stress that upregulates the production of PR-5 proteins in the plant. Also, the mature fusion protein showed no effect on spore germination of the fungi (data not shown). Nevertheless, most of the pathogens tested did not exhibit any growth inhibition by rTLP; *in-vitro* assay shows that it has some effect on mycelial growth of the fungi tested. After 24 h of further incubation, discs were overgrown by the fungal mycelia. This further confirms its static nature, but not lethal.

**Figure 2A, 2B:**
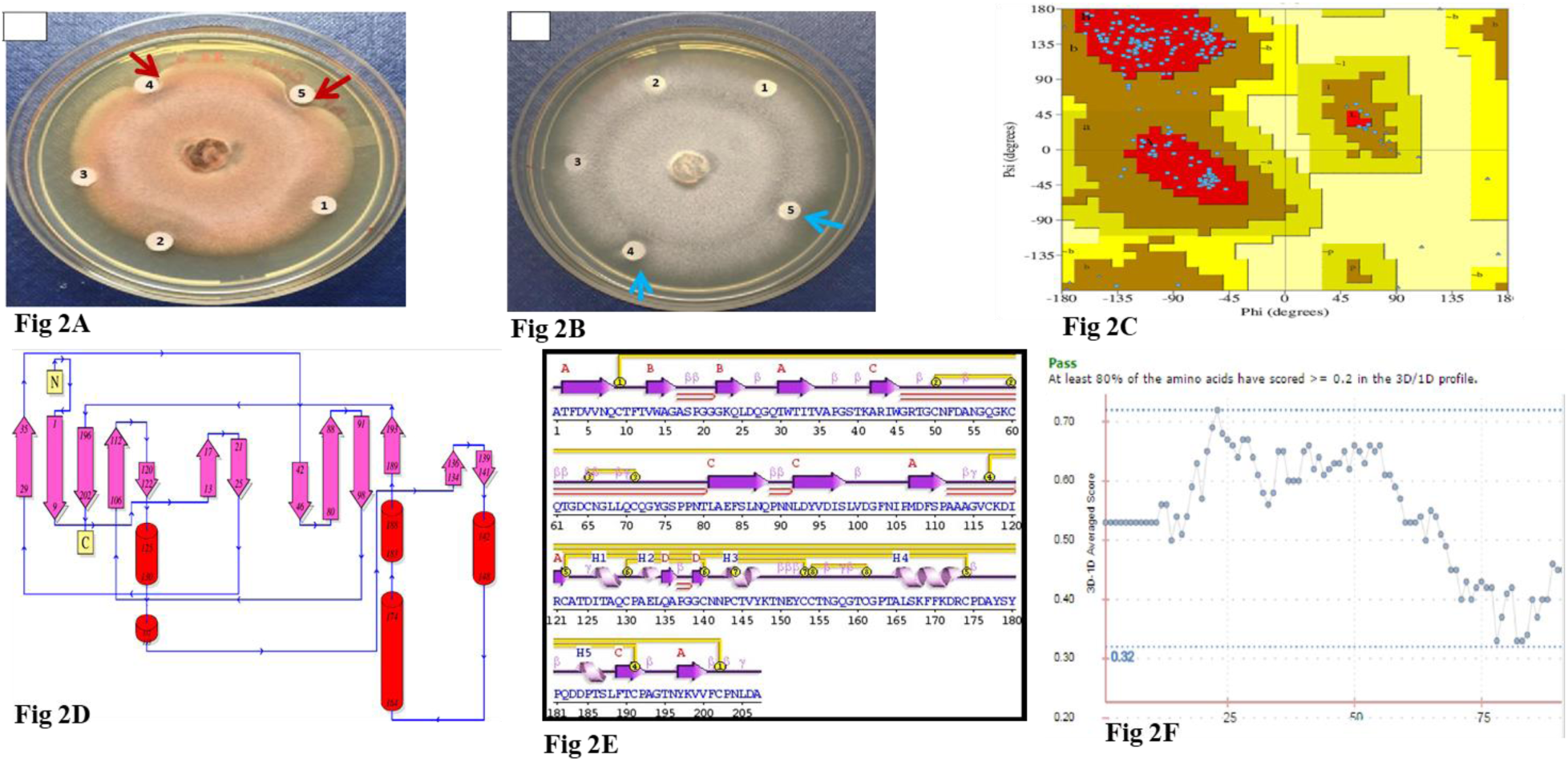
Bioassay for mycelial growth inhibition activity by heterologous expressed and refolded poly-histidine-tagged mature acidic TLP (rTLP). **Fig. 2A and B** Representative Petri dishes of disc diffusion assay with fungal species, *Alternaria alternata* and *Geotrichum candidum*. Filter discs of 6.0 mm diameter saturated with 50 μL of test protein solution were placed adjacent to actively growing fungal mycelia. 20 μg of BSA was used as negative control. Inhibitory effects of rTLP against the fungi are observed as area lacking the growth of mycelia. The numerical 1, 2, 3, 4, and 5 indicates various concentrations of rTLP used for testing such as 12.5 μg, 25 μg, 50 μg and 100 μg of protein. A dose of 20 μg of BSA was used as negative control **Figure. 2C:** Ramachandran plot statistics for computer-generated model of rTLP. The figure shows amino acid residues in most favourable regions (A, B, L) 85.4 %, additionally allowed regions (a, b, l, p) 14.6 %, generously allowed regions (∼a, ∼b, ∼l, ∼p) and disallowed regions. **Figure. 2D:** Topology diagram of rTLP structure. The figure represents the presence of 13 well conserved β-strand that runs anti-parellel to each other where 11 strands belong to domain 1 and the other 2 strands belong to domain III; Five helices that belongs to domain II of rTLP. The predicted secondary structure is in accordance with the defined structure of rTLP. **Figure. 2E:** Secondary structure of rTLP predicted using PROTIF-scan: *Sapodilla* acidic TLP comprises of 13 β-strands; 5-helices; and conserved disulfide linkages are represented by yellow lines. **Figure. 2F:** A verify 3D-plot for validation of the generated model. The plot is generated by comparing the 3D structure with the primary sequences of the protein profile is created based on the local environment of the residues.

### Structural bioinformatics rTLP

The rTLP sequences were subjected to bioinformatics study wherein the PROCHECK analysis, showed 85.4 % in most favoured regions, 14.6 % in additional allowed regions, no residues in generously allowed regions and disallowed regions (Fig. 2C, D). The assessment of the overall structure based on the plot characteristics reveals that the built model is good with acceptable stereo-chemical quality.

The ‘proportion secondary structure’ of rTLP shows presence of 13 β-strands where 11 β-strands belong to domain I and the other two β-strands fit into domain II of the protein. The presence of 16 cysteine residues and the disulfide bridging are clearly depicted in the Fig. 2E, F. The diagram also shows the presence of less α-helix content. These structural details are in accordance with the reported structural traits of TLP’s. The rTLP structures were predicted and shown in Fig. 3A, B, C.

**Figure 3A:**
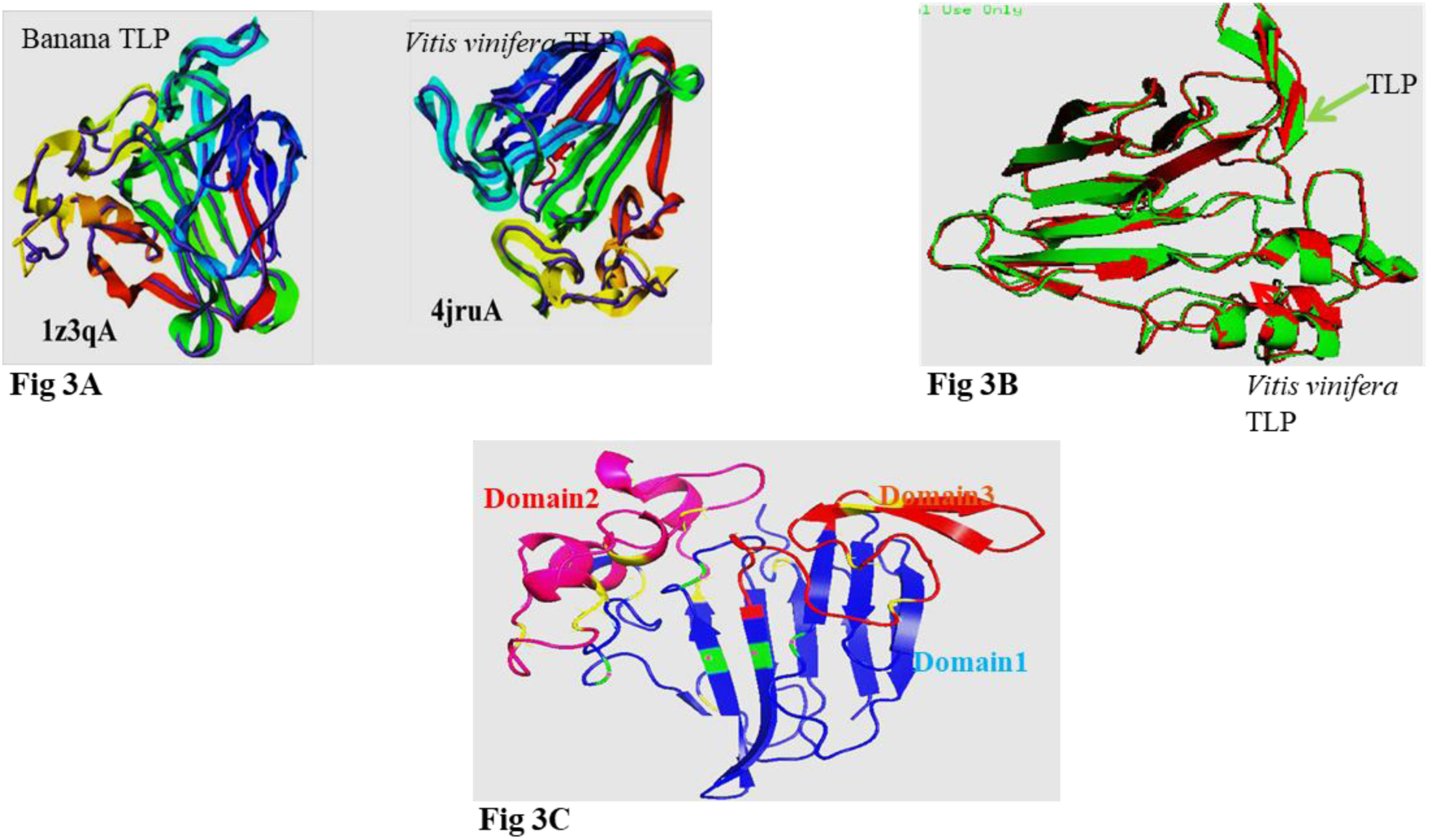
PDB structures of top two templates used by I-TASSER for model generation. 1z3qA represents PDB ID of Banana TLP; 4jruA represents PDB ID of *Vitis vinifera* TLP. The templates are represented in rainbow-colored cartoons and predicted model of TLP is represented ribbon structures. **Figure. 3B:** A superimposed representation of I-TASSER generated model of TLP(green) with the structurally closest PDB hit *Vitis vinifera* TLP(red) (PDB:4jru) **Figure. 3C:** A predicted three dimensional homology model of *Sapodilla* acidic TLP generated by I-TASSER. The model shows three domains of TLP generated by I-TASSER. The central β-barrel domain (domain 1) forming the hydrophobic core of the molecule is shown in blue. The domain II formed by α-helices is shown in pink. The domain III formed by two short β-strands are shown in red. The disulfide bridges are shown in yellow. The residues of REDDD cleft are shown in green.

### Immuno-detection of rTLP variants

The rTLP variants were studied for their cross-reactivity with antibodies generated from TLP protein. The purified mutants were resolved by SDS-PAGE and transferred onto PVDF membrane followed by immune-detection with anti-TLP antibodies. Both the variants showed a positive reaction to polyclonal antibodies indicating the cross-reactivity (Supplementary Fig. 1A, B, C). Supplementary Figure. 1A, B represent purification profile of rTLP variants and Figure 1C represents western blot. This result suggests that the epitope regions of the variants post-mutation remained unaltered. The western blot profile also indicates the absence of dimer in both the variants. This could be further speculated to understand the role of the mutated residues in the formation of dimers of the variant proteins (supplementary data; Fig. 1 Deletion mutant purification, Fig.1B: Site specific mutant purification and Fig. 1C. Western blotting of rTLP, Del and SDM mutants of rTLP).

## Discussion

Structurally and catalytically important amino acid residues are thought to be conserved in protein evolution. TLPs from different plant sources have been purified and studied extensively due to their allergy and antifungal properties. TLPs are unique members of the PR family exhibiting many distinct biological functions in the plant system. Two properties of TLPs have been extensively studied: one, the antifungal activity and the other its enzymatic activity that catalyses the hydrolysis of the β-1, 3-glucans. The mechanism by which the TLPs counter-attack fungal pathogens has not been fully realized. However, initially, it was speculated that the hydrophobic core of the molecule accommodates itself into the cell walls of fungal pathogens resulting in the compromised cell wall and finally leading to fungal death [28].

The results indicate that rTLP shows sensitivity to two fungal species. The growth of *Alternaria alternata* and *Geotrichum candidum* was very weakly inhibited by rTLP (19.8 % and 16 % inhibition at 100 μg concentrations of test protein, respectively) at a relatively high concentration of the protein. Hence, rTLP may possess fungistatic activity rather than being a potent fungicide for the above-tested pathogens.

Of late, revelations on endo-β-1, 3-glucanase activity of TLPs have substantiated the theories about TLPs exerting the antifungal effect at the level of fungal cell walls. The presence of inter-domain electronegative cleft in more or less all the TLPs with antifungal activity supports the above observations. It is also noteworthy that TLPs lacking enzymatic activity also shows resistance to the growth and propagation of fungal species. Different isoforms of TLPs reported from the same plant species are known to exhibit antifungal as well as enzymatic activity with different intensity [29]. Although the members of PR-5 members show remarkable similarity to each other by both sequence and structure, functionally the members vary in different aspects. The conservation of function in different members of the PR-5 family is still not well-understood. Many studies have examined the conservation of structure in evolutionary perspective, but not through site-directed mutagenesis approach. Hence, an effort has been made to understand the relationship between the conserved structural scaffolds and their functional variations.

In the present study, the molecular model of TLP was generated using the I-TASSER tool. I-TASSER generated model with grape TLP as a structural homolog template was taken for further validations and analysis of structural features. The predicted structure of TLP was in agreement with structural descriptions of several other TLPs (Fig. 3A, B, C). The model was further validated to be energetically stable by PROCHECK, ERRAT and Verify 3D online validation programs. Further docking studies with the generated model which reveal the amino acid side chains involved in enzyme catalysis and antifungal property could be of increasing interest.

TLPs are mainly assigned to their conformation stabilized by eight disulfide linkages that render the protein its resistance to thermal, chemical (e.g., urea) and proteolytic degradation [30]. The polypeptide folds into three distinct domains that are highly conserved. It is observed that domain I of the protein forms the hydrophobic core of the molecule. The folding of a protein is mediated by the establishment of an intricate disulfide network, which forms three distinct domains with high hydrophobicity [31]. The disulfide linkage is intrinsically a local phenomenon for each polypeptide. Apparently, the appearance of disulfide bridging is more pronounced in small proteins than in the large proteins. This indicates that disulfide bridging in smaller proteins is likely to have an important role in stabilization mainly than in larger counterparts. It has been proposed that upstream half-cysteines have downstream neighbours and conversely, downstream cysteines prefer upstream residues for linkages, a fact consistent with the role of cysteines as ‘structure-keepers’. It is important to assess the relevance of particular disulfide bridges in structural stability and functionality [32].

Similarly, substantiating the above finding, in *Sapodilla* TLP, the C^9^ in the N-terminal region is paired with the C-terminal region C^202^ forming the outer loop of the protein, as it is in other TLPs. The C-terminus of TLP (-VVFCP region) is known to be conserved and harbours a cysteine residue involved in disulfide pairing. The preliminary understanding of the importance of the C-terminal tail in functionality has been made by generating a C-terminally truncated protein that lacks a nonapeptide (-VVFCPNLDA).

The mutant was cloned successfully into pET-23d ^(+)^ vector to obtain pET-23d ^(+)^ -ΔCrTLP. The expressed protein was refolded by solubilizing the inclusion bodies in 6.0 M urea. The IBs were solubilized in 6.0 M urea solution, whereas native rTLP required 8.0 M urea for its denaturation suggesting that the C-terminally truncated protein could be more susceptible to chemical denaturation. This further shows that the C-terminal region of the protein is an important component for the stability and folding of the protein. The refolded and purified protein was assayed qualitatively for the enzymatic and antifungal activity. The enzymatic activity of the C-terminal truncated protein was found to be dramatically decreased where there was a complete loss of the antifungal activity. The crystal structure of several TLPs have been resolved so far to elucidate the mechanism of glycosidic bond hydrolysis in polysaccharide substrates. The conformational electronegative cleft formed by a cluster of five residues (REDDD) is known to be conserved and responsible for the β-1, 3-glucanase activity of TLP. As shown in other TLPs, e.g., NP-24, the polysaccharide chain is fitted into the inter-domain cleft where the substrate is surrounded by Arg, Glu, Asp, and other aromatic amino acid residues. In particular, Glu^84^, Asp^97^, and Asp^102^ are known to form hydrogen bonding with the substrate. Based on the three-dimensional structure, and the multiple sequence alignment studies, in *Sapodilla* TLP, the residues Glu^83^, Asp^96^, and Asp^101^ are expected to form the catalytic centre required for the endo-β-1, 3-glucanase activity of rTLP.

Hence, in an attempt to study the role of Asp^96^ residue in the catalysis of glucan-hydrolyzing mechanism, alteration of ^D^96^R^ was carried out by site-directed mutagenesis. The mutant ^D^96^R^rTLP fused with poly-histidine-tag at the C-terminal end was successfully cloned and expressed as inclusion bodies in BL21 (DE3) *E. coli* cells. The mutant protein was subjected to on-column refolding in the presence of folding enhancers (GSH: GSSG) and purified to homogeneity by using metal-ion chromatography. The β-1, 3-glucanase activity of the mutant was qualitatively assayed by zymography with laminarin as a substrate. The results show that the replacement of ^D^96^R^ drastically affects the enzymatic activity of the protein, which falls below the minimum level of detection.

The sTLPs (small-TLPs), a subset of thaumatin-like family proteins are small proteins of around 16 kDa sharing high structural similarity with thaumatins. These proteins contain only five disulfide bonds of the eight linkages in TLPs. The internal deletion of ∼58 amino acid residues is attributed to the loss of three disulfide linkages. However, the disulfide bond corresponding to the C_N_ (N-terminal cysteine) and C_C_ (C-terminal cysteine) is shown to be preserved during evolution. This could plausibly explain the importance of C^9^-C^202^ disulfide loop in maintaining the stability and biological activity of rTLP. Hence, the present study suggests that both C-terminal nonapeptide and ^D^96 residues contribute to the functional activity of rTLP.

The assessment of the overall structure based on the plot characteristics reveals that the built model is good with acceptable stereo-chemical quality (Fig. 2C, D). The Verify 3D profile for the predicted structure shows that 80 % of the residues had an average score over 0.2. Thus the predicted structure for rTLP was considered reliable (Fig. 2E, F). The overall quality factor of the predicted model was found to be 93 %. The good models generally produces values above 91 %. Hence, this program substantiates the fact that the model built was consistent and reliable.

Heterologous expression of protein has been an approach of choice for producing polypeptides that are available in limited quantities from natural sources. The results described here suggest that the gene encoding *Sapodilla* acidic TLP have been successfully cloned and expressed in *E. coli*. The overexpression of rTLP was standardized for refolding and purification. However, the refolding of proteins in its functionally active and soluble form is still a bottleneck. TLP is a disulfide-rich protein has offered an exciting challenge for optimizing the convenient and ideal refolding method. This study describes three different methods of refolding and purification. They determine the best method of choice for functional expression and purification of rTLP. It is likewise shown that the presence of reduced and oxidized glutathione is critical to aid native folding of disulfide-rich proteins. The chromatographic refolding procedure in the presence of redox-folding enhancers has shown promising results. Overall, downstream processing of recombinant TLPs offers an excellent platform to obtain recombinant proteins of high purity and sufficient yield compared to the native proteins for investigation of the structure-function relationships.

The recombinant expression and selected protein engineering of acidic TLP from *Sapodilla* plum have been a unique prototype to study the structure-function relationship. This study has been a preliminary attempt to assess the role of cysteine in the C-terminal region. And also aspartic acid residue (^D^96) in the REDDD electronegative cleft in the functional properties of rTLP by mutagenesis approach. The drastic loss in the enzymatic activity of the mutated protein has suggested the participation of these residues in the structure and function of TLP. This investigation suggests that the disulfide linkage (C^9^-C^202^) forming the outer loop of the protein is responsible for keeping the active site residues in place by maintaining the stability of the protein. The current study may be extended to understand the role of other disulfide determinants in overall stability and retention of functional attributes.

The refolded mature rTLP was assayed for its antifungal property. The mycelial growth of *A. alternata* and *G. candidum* was observed to be slightly inhibited by this protein, thus displaying its weak antifungal activity against two phyto-pathogenic fungi. LocTree3 suggested the protein be targeted into extracellular space which also substantiates the above experimental results, indicating that acidic isoform of rTLP in *Sapodilla* might be involved in the basal defence response of the host.

## Supporting information

Purification of TLP variants

## Acknowledgements

CG acknowledges CSIR-CFTRI and AcSIR for facilities and financial support.

We acknowledge CSIR-CFTRI for granting a fellowship during PhD to CTG, and also for providing infrastructural facilities.

---

**Supplementary Figure 1A, B, C:**
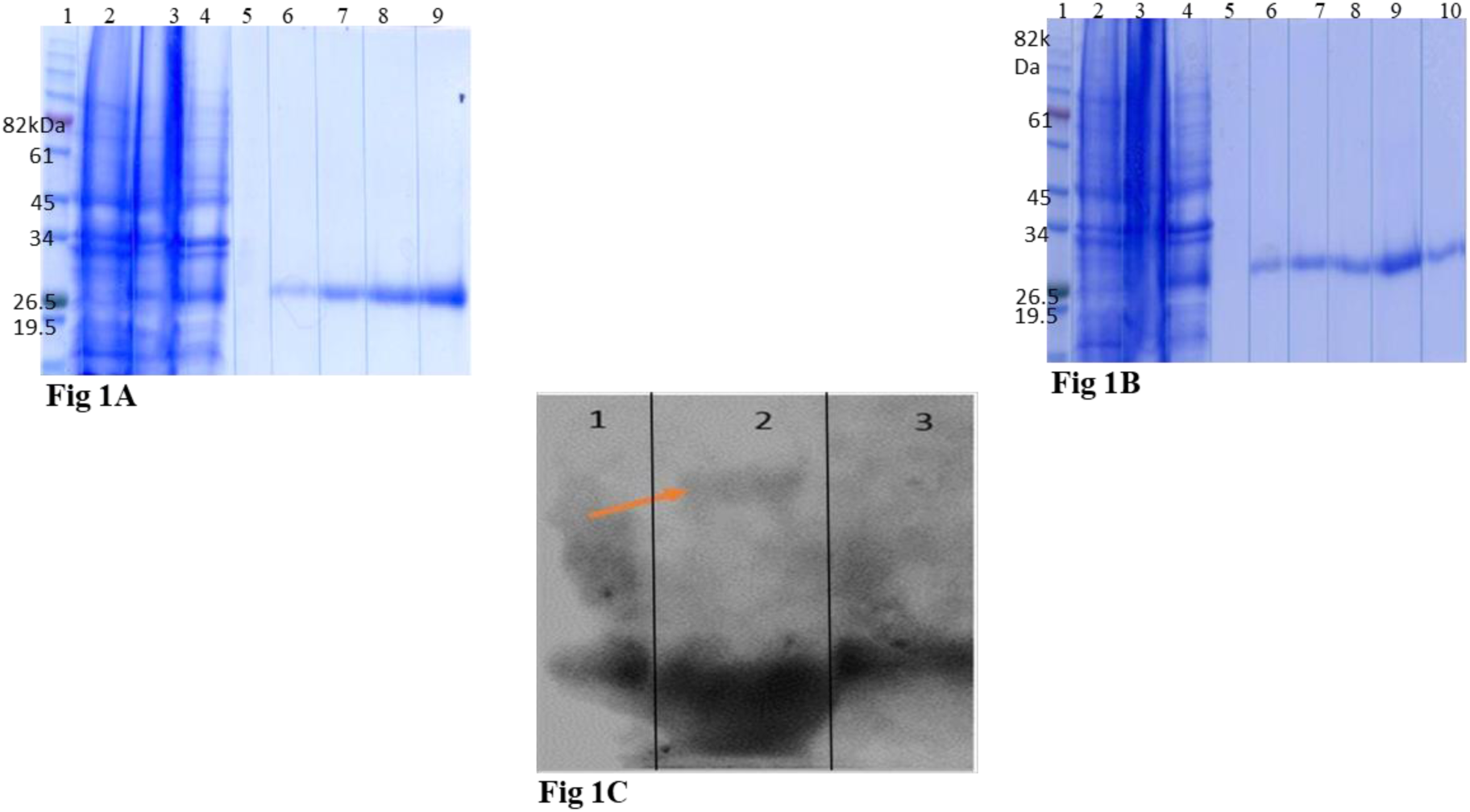
**Figure 1A:** Expression and metal-ion affinity chromatography profile of ΔCrTLP1 truncated protein expressed as C-terminal his-tagged protein. SDS-PAGE profile of selected elution fractions; Lane 1: Protein marker; Lane 2: Un-induced cell lysate; Lane 3: Induced cell lysate; Lane 4: Sediment of sonicated cell lysate; Lane 5: Wash with 50 mM imidazole; Lanes 6-9: Eluted fractions with 0.3 M imidazole. **Figure 1B:** Expression and metal-ion affinity chromatography profile of ^D^96^R^rTLP mutant expressed as C-terminal his-tagged protein. (A) The elution profile of mutant protein with 0.3 M imidazole; (B) SDS-PAGE profile of selected elution fractions; Lane 1: Protein marker; Lane 2: Un-induced cell lysate; Lane 3: Induced cell lysate; Lane 4: Sediment of sonicated cell lysate; Lane 5: Wash with 50 mM imidazole; Lanes 6-10: Eluted fractions with 0.30 M imidazole. **Figure 1C:** Immuno detection of variants with anti-TLP antibodies. Lane 1: Δ CrTLP; Lane 2: rTLP1; Lane 3: ^D^96^R^ rTLP. An arrow in Lane 2 indicates the presence of homodimer of rTLP wild type protein whereas the formation of dimer is not observed in both the variants Lane 1, and 3.

