## Supplementary figures and images for "*Sapodilla* rTLP exists as a monomer, dimer with β-1, 3-glucanase and antifungal activity"

### Purification of TLP variants

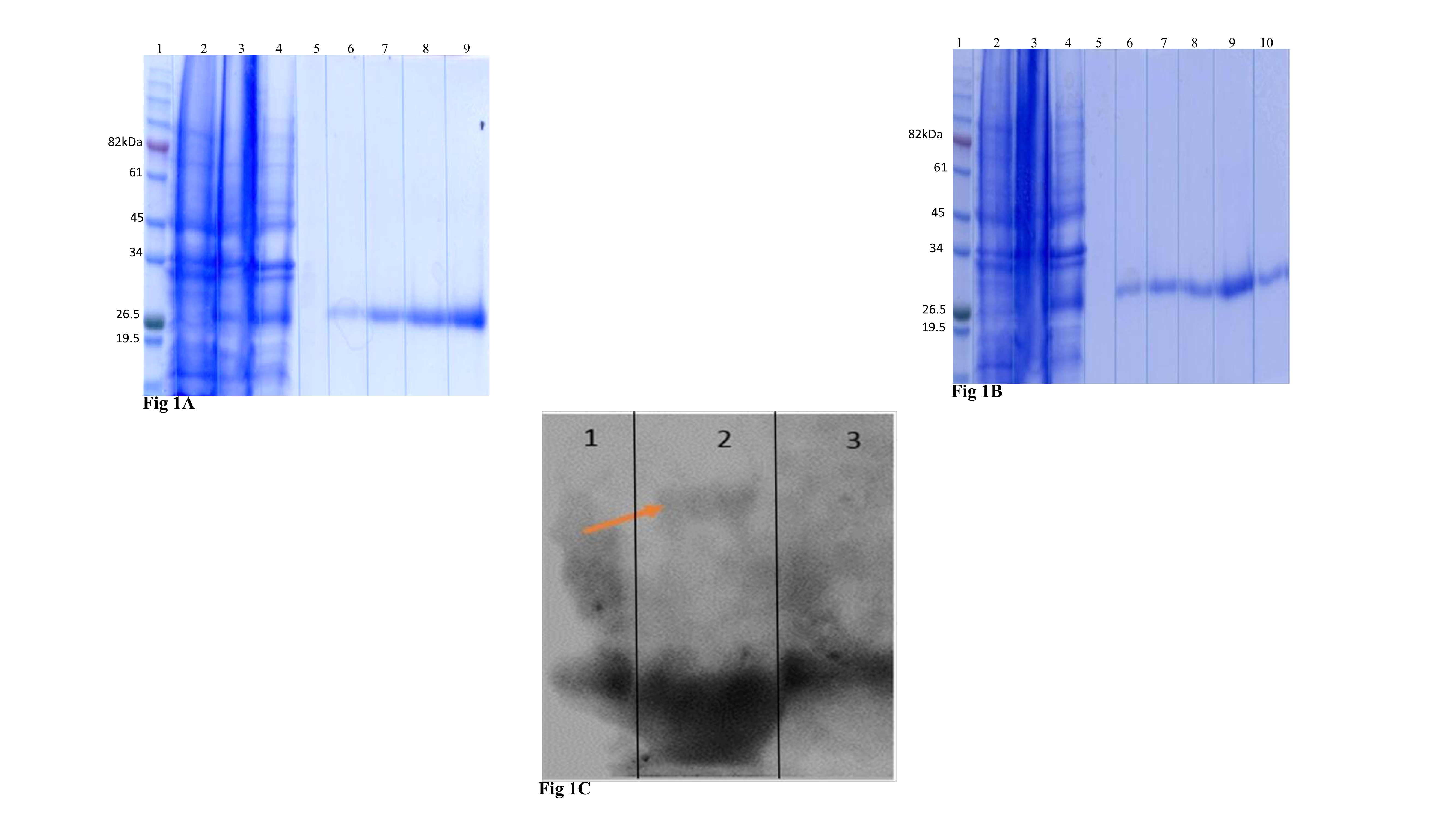
